# Environment-driven reprogramming of gamete DNA methylation occurs during maturation and is transmitted intergenerationally in salmon

**DOI:** 10.1101/2020.11.25.396069

**Authors:** Kyle Wellband, David Roth, Tommi Linnansaari, R. Allen Curry, Louis Bernatchez

## Abstract

An epigenetic basis for transgenerational plasticity is widely theorized, but convincing empirical support is limited by taxa-specific differences in the presence and role of epigenetic mechanisms. In teleost fishes, DNA methylation generally does not undergo extensive reprogramming and has been linked with environmentally-induced intergenerational effects, but solely in the context of early life environmental differences. Using whole genome bisulfite sequencing, we demonstrate that differential methylation of sperm occurs in response to captivity during the maturation of Atlantic Salmon (*Salmo salar*), a species of major economic and conservation significance. We show that adult captive exposure further induces differential methylation in an F1 generation that is associated with fitness-related phenotypic differences. Genes targeted with differential methylation were consistent with genes differential methylated in other salmonid fishes experiencing early-life hatchery rearing, as well as genes under selection in domesticated species. Our results support a mechanism of transgenerational plasticity mediated by intergenerational inheritance of DNA methylation acquired late in life for salmon. To our knowledge, this is the first-time environmental variation experienced later in life has been directly demonstrated to influence gamete DNA methylation in fish.

## Introduction

The inheritance of environmentally-induced epigenetic variation (e.g. DNA methylation, chromatin structure, small RNAs) has been proposed as a mechanism facilitating transgenerational plasticity (Richards 2006; Bell and Hellmann 2019). As a chemical modification of nucleotide bases, DNA methylation has a clear mechanism for multigenerational transfer, however our current understanding of its role in intergenerational (i.e. parent-offspring transfer) or transgenerational (i.e. multi-generational transfer) epigenetic inheritance in animals is hindered by a lack of data from a wider diversity of organisms and thus evidence remains limited and controversial (Heard and Martienssen 2014; Horsthemke 2018; Skvortsova *et al.* 2018). In mammals, DNA methylation undergoes extensive reprogramming (de-methylation and re-methylation) following fertilization and then again when germ-line tissue differentiates from somatic tissue (Heard and Martienssen 2014). This generally limits the potential for intergenerational inheritance as methylated sites must escape two rounds of re-patterning in each generation. Other well-studied traditional animal models (i.e. worms and flies) lack methylation and thus provide no relevant insight. However, a fish model, specifically zebrafish (*Danio rerio*), does not exhibit global erasures and reprogramming of methylation neither following fertilization where the maternal pattern is reprogrammed to match the paternal pattern (Jiang *et al.* 2013; Potok *et al.* 2013) nor during germline differentiation (Ortega-Recalde *et al.* 2019; Skvortsova *et al.* 2019) suggesting a greater potential for DNA methylation-mediated epigenetic inheritance in this species and possibly teleost fishes more generally (however see: Wang and Bhandari 2019).

Environmentally-induced DNA methylation variation in a wider selection of teleost fishes has been reported as a result of differences in early developmental environments (Le Luyer *et al.* 2017; Artemov *et al.* 2017; Metzger and Schulte 2017; Gavery *et al.* 2018). These signals stably persist until adulthood (Metzger and Schulte 2017; Leitwein *et al.* 2021), occur in germ cells (Gavery *et al.* 2018; Rodriguez Barreto *et al.* 2019), and a growing body of literature demonstrates that intergenerational transmission can occur (Ryu *et al.* 2018; Rodriguez Barreto *et al.* 2019; Berbel-Filho *et al.* 2020; Heckwolf *et al.* 2020), thus supporting an overall mechanism of DNA methylation-mediated intergenerational epigenetic inheritance for this clade.

The timing for germ-line incorporation of environmentally-induced epigenetic variation is historically believed to be limited to early developmental stages as a result of the separation of the germline and soma, or the so-called “Weismann Barrier” (Monaghan and Metcalfe 2019). Once germline epigenetic reprogramming is completed, this barrier limits somatic influence on the germline and thus, would theoretically be expected to prevent environmentally-derived epigenetic changes from being incorporated into gametes and passed to future generations. Yet, emerging evidence suggests that the barrier is more permeable than previously thought (Eaton *et al.* 2015), with evidence is for small RNA mediated epigenetic inheritance (Sciamanna *et al.* 2019; Duempelmann *et al.* 2020) and at least one example of differential methylation in mice (Dias and Ressler 2014). However, wide taxonomic evidence for these phenomena are lacking and this makes it difficult to assess the potential for intergenerational transmission of environmentally-induced DNA methylation variation acquired after early developmental stages.

Salmonid fish hatcheries provide relevant systems in which to test hypotheses regarding intergenerational epigenetic inheritance. Hatcheries have been used for decades to enhance, supplement, or recover salmonid fish populations (Naish *et al.* 2007), but have negative consequences for the fitness of hatchery-reared fish (Araki *et al.* 2008; Christie *et al.* 2014; O’Sullivan *et al.* 2020) presumably due to domestication effects. Numerous studies have failed to demonstrate significant genetic difference between hatchery-origin and natural-origin (wild) salmon (Christie *et al.* 2016; Le Luyer *et al.* 2017; Gavery *et al.* 2018) despite documented pronounced differences in gene expression (Christie *et al.* 2016). In contrast, DNA methylation divergence has been reported between hatchery-origin and wild salmon for several species (Le Luyer *et al.* 2017; Gavery *et al.* 2018, 2019; Rodriguez Barreto *et al.* 2019). Evidence for intergenerational effects has recently emerged (Rodriguez Barreto *et al.* 2019), although solely in the context of early developmental hatchery exposure where salmon are artificially reproduced and the offspring reared in the hatchery environment for a period of time before release into the natural environment. Assuming the Weismann Barrier exists in salmon, environmentally induced epigenetic differences would be expected to arise shortly following fertilization and before the differentiation of primordial germ cells.

Alternative hatchery rearing techniques for salmon, including juvenile (here: smolt) to adult supplementation (hereafter SAS; as per Fraser 2016) and live gene-banks, are increasingly being applied to conserve and recover the most critically endangered salmon populations (O’Reilly and Doyle 2007; Stark *et al.* 2014). In these conservation strategies, wild-born juvenile fish are collected from the wild, reared to adulthood in a hatchery environment, and at the onset of maturation they are again released to the wild to spawn naturally. These strategies are believed to reduce the risks to wild populations that are associated with the use of hatchery technology by allowing natal stream imprinting by juveniles as well as homing, natural mate-choice, and spawning site selection by adults. While these approaches show promise for demographic recovery of some populations (Berejikian and Van Doornik 2018), fitness-related differences between SAS and wild salmon have also been documented (Berejikian *et al.* 2001, 2005). In these contexts, it is unclear whether environmentally derived epigenetic modifications are a plausible explanation for the observed phenotypic effects given that germ-line DNA methylation patterns, assuming an impermeable Weismann Barrier, should have been determined well before fish experienced a captive environment. As such, we currently lack knowledge of the potential for epigenetically-mediated intergenerational effects that may result in heritable declines in fitness in these contexts. These systems also provide opportunities to test the potential for intergenerational transmission of late-life, environmentally-induced, DNA methylation variation.

In this context, the goal of our study was to test the hypothesis that environmental differences experienced by adults during maturation alter DNA methylation of gametes. We further test the hypothesis that adult rearing environments influence offspring methylation patterns and that these differences influence offspring phenotypes.

## Methods

Atlantic Salmon (*Salmo salar*) juveniles (smolts) (Figure 1A) were collected using a rotary screw trap from May to June 2015 from the mainstem of the Northwest Miramichi River, New Brunswick, Canada near the mouth of Trout Brook (Figure 1B). They were transported to the Miramichi Salmon Conservation Centre (MSCC, South Esk, NB; Figure 1C) where they were held in tanks with natural photoperiod until 2017 (i.e. smolt-to-adult supplementation or SAS). SAS fish were initially fed frozen shrimp and gradually weaned onto standard hatchery pellet food over the course of a couple months. Natural origin (wild) adult Atlantic Salmon returning to spawn in the Miramichi River to spawn were collected by seining in September of 2017 and 2018 as a part of regular brood-stock collection program by the Miramichi Salmon Association staff from selected pool habitats in the upper reaches of a major branch (the Little Southwest Miramichi River) of the Northwest Miramichi River (Figure 1E). Adult fish were transferred to MSCC and held in tanks for up to three weeks.

**Figure 1:**
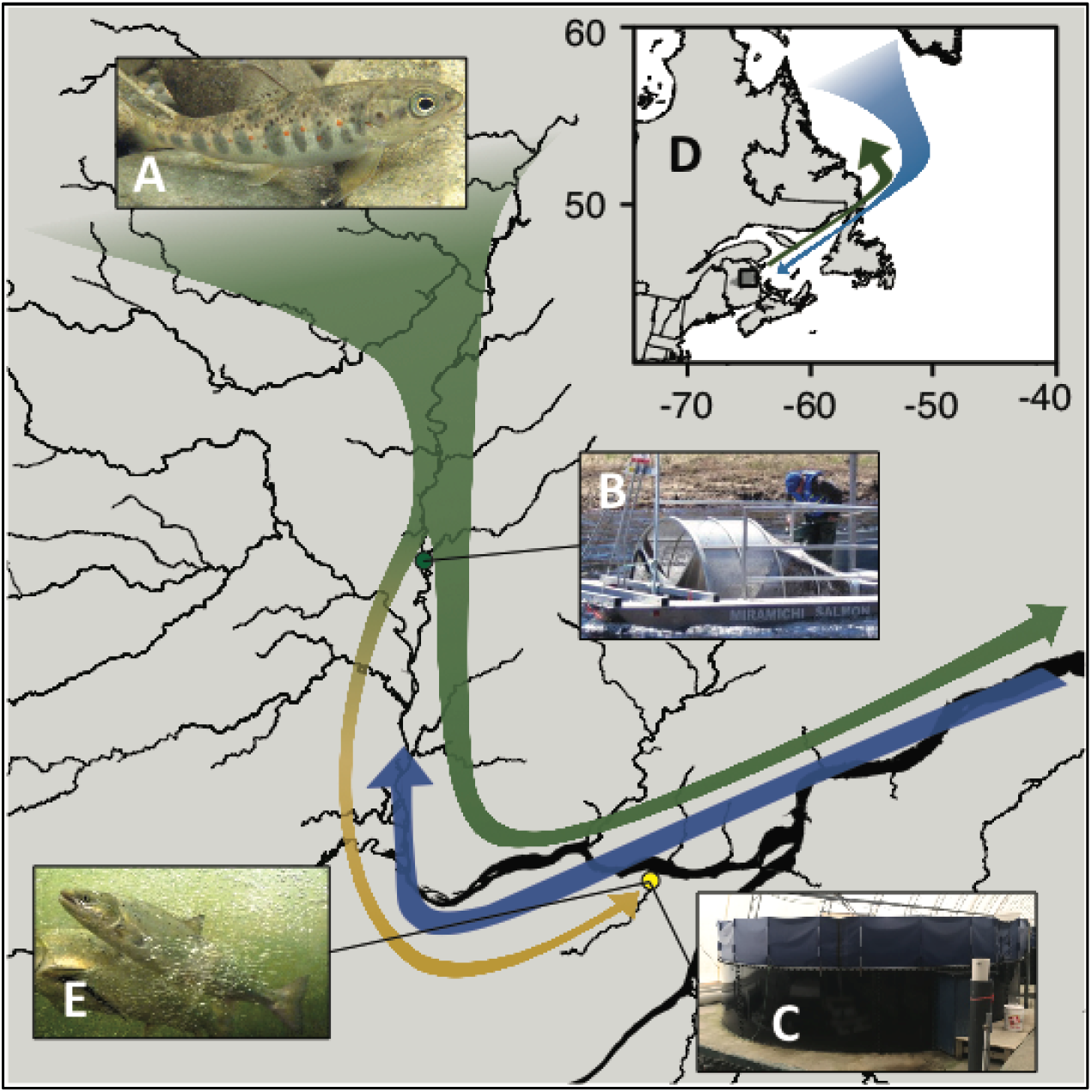
Smolt-to-adult supplementation in the Northwest Miramichi River. Natural origin (wild) juvenile salmon (A) are captured during their migration to the ocean (B) and reared in captivity until adulthood (SAS) at the Miramichi Salmon Conservation Centre (C). Wild salmon continue their marine migration spending 1-3 years feeding in the Labrador Sea (D) before returning to the Miramichi River to spawn. Wild salmon from the same cohort as SAS salmon were captured in various headwater pool habitats during their return migration (E).

In 2017, both SAS and wild adult fish were haphazardly netted from holding tanks, gametes were collected, and eggs were artificially fertilized to create pure-type breeding crosses (SAS x SAS and wild x wild; N = 8 crosses each). Fertilized eggs were incubated in flow-through troughs until first feeding after which offspring from multiple families were mixed and transferred to 3m x 3m square tanks and fed ad libitum on hatchery pellet food until they reached a size of approximately 1.5 g. Juvenile offspring fish were netted haphazardly from the tanks, euthanized in an overdose solution of eugenol (Sigma-Aldrich Canada, Oakville, ON), weight and length measurements taken, and liver tissue was dissected and preserved in RNAlater. Juvenile samples were later genotyped with a panel of 188 SNPs (KASP SNP assays; LGC Biosciences, Beverley, MA, USA) and assigned to their family of origin using COLONY v2.0.6.5 (Jones and Wang 2010). In 2018, sperm samples were collected from eight wild and eight SAS males in 2 mL tubes and stored on ice for between 2 – 6 hours. Sperm (250 μL) was centrifuged at 7000 rpm for 10 min, the supernatant discarded, and isolated sperm cells preserved in 1.5 mL of RNAlater.

Genomic DNA was extracted from sperm and liver tissue using a salt-based extraction protocol (Aljanabi and Martinez 1997). Whole-genome bisulfite sequencing libraries were prepared at the McGill University - Génome Québec Innovation Centre using NEBNext Ultra II DNA library prep kits. Each library was sequenced to an expected depth of 15X using 150 bp paired-end sequencing on a half lane of the Illumina HiSeqX platform (Table S1). Raw sequencing reads were trimmed using fastp v0.19.9 (Chen *et al.* 2018). Trimmed reads were aligned to the Atlantic Salmon genome (ICSASG v2; NCBI RefSeq: GCF_000233375.1; Lien *et al.* 2016) with bwameth v0.2.2 (Pedersen *et al.* 2014). Duplicate reads were removed using the MarkDuplicates tool from the Picard Toolkit v2.23.1 (Broad Institute 2019). Methylation statistics for all CpG dinucleotides were compiled using MethylDackel v0.4.0 (Ryan 2019) excluding reads with less than 15 bases possessing a mappability greater than 0.01 as calculated by Bismap v1.1.1 (Karimzadeh *et al.* 2018) for 150 bp reads. Detailed scripts including parameter values are available on github: https://github.com/kylewellband/bwa-meth_pipeline.git.

To characterize C-T polymorphisms that could bias methylation estimates, we combined equal proportions of DNA from all individuals (N = 36) and sequenced them as a pool in one lane of an Illumina HiSeqX. Raw sequencing reads were trimmed using fastp v0.19.9 and then aligned to the Atlantic Salmon genome with bwa mem (Li 2013). Duplicate reads were removed using MarkDuplicates and overlapping 3’ ends of paired reads were clipped using the clipOverlap function of BamUtil v1.0.14 (Jun *et al.* 2015). We called SNPs using a frequency based approach in freebayes v1.3.1 (Garrison and Marth 2012) that required variant sites to be covered by a minimum of 10 reads and have a minimum of two reads supporting the alternate allele. We retained both C-T and A-G (i.e. C-T on the minus strand) polymorphisms and removed these sites from the methylation results using bedtools v2.26.0 (Quinlan and Hall 2010).

We quantified differential methylation at CpG sites covered by at least one read in all samples where we additionally required a minimum of five reads and a maximum of 20 reads (approximately 99.9^th^ quantile) for at least 12 of 16 juveniles or eight of 12 adults. The sequencing performance for four adult samples (i.e. two HiSeqX lanes) representing two SAS and two wild fish was poor and thus these samples were excluded to reduce biases in methylation estimates due to low coverage for these samples. The minimum coverage filter ensured that differences in methylation were not due to spurious differences in coverage between groups and the maximum coverage filter removed highly repetitive regions where the confidence in mapping accuracy is low. Differential methylation of CpG cytosines (DMC) was determined using beta-binomial models implemented in the DSS v2.32.0 package (Feng *et al.* 2014) in the R statistical environment v3.6.1 (R Core Team 2019). Methylation levels were first smoothed using a window size of 500 bp and models were fit with group-specific dispersion estimates as implemented in DSS. False discovery rates were calculated according to Benjamini and Hochberg (1995). Differentially methylated regions (DMRs) were determined based on attributes of DMCs, where regions were required to be a minimum of 50 bp long, have > 3 CpGs, and greater than 50% of the CpG sites with a p-value < 0.001. Due to the large number of small contigs in the Atlantic Salmon genome (i.e. >230,000; ICSASG v2; NCBI RefSeq: GCF_000233375.1; Lien *et al.* 2016), we restricted our analyses to the 29 full-length chromosomes and contigs larger than 10 kb in length (>96% of the un-gapped length of the genome).

To determine potential functional consequences of methylation differences we used bedtools v2.26.0 (Quinlan and Hall 2010) to identify gene features associated with DMRs. NCBI RefSeq gene annotation information for the salmon genome was retrieved and genes were associated with differential methylation if any DMRs were located within 5000 bp of a coding region consistent with previous work in salmonids (Le Luyer *et al.* 2017). Gene ontology information for annotation genes was obtained from the Ssal.RefSeq.db v1.3 (https://gitlab.com/cigene/R/Ssa.RefSeq.db; accessed: June 22, 2020) R package. We tested for enrichment of biological functions for genes associated with DMRs using Fisher’s Exact Tests and the ‘weight01’ (Alexa *et al.* 2006) as implemented in the TopGO v.2.38.1 package (Alexa and Rahnenfuhrer 2019).

We used a network-based approach to investigate associations of correlated methylation signatures with juvenile phenotypes. We first summarized methylation in non-overlapping 100 bp windows across the genome. Windows required a minimum of three CpG sites and we only retained windows with among-sample variances greater than 0.05 to reduce computational burden when constructing the network (N = 59,803 windows). We calculated connectivity between all pairs of regions using the bi-weight midcorrelation raised to the power of 18 to approximate a scale-free network as implemented in the WGCNA package in R (Langfelder and Horvath 2008). Modules of correlated methylation signatures were inferred using hierarchical clustering of the topological overlap dissimilarity matrix and a dynamic tree-cutting algorithm. The modules were constructed using a block-wise approach with a maximum of 30,000 regions allowed in each block and all blocks were then merged to form the final modules. The association of methylation modules with phenotypes was assessed by correlating module eigenvectors scores (the first axis of a principal component analysis conducted on all the regions within each module) with phenotypic values for each individual using the bi-weight midcorrelation. Module-traits correlations with p-values < 0.05 were retained for further analysis. For each significant module-trait correlation we assessed whether the any regions within the module overlapped with previously identified DMRs found between SAS and wild fish. We assessed the statistical significance of these associations using a resampling procedure to compare the number of DMR overlaps with those based upon a random draw of regions for each module. We tested for differences in average module membership (region correlation with module eigenregion) and gene significance (region correlation with phenotype) between DMR-overlapping and non-DMR-overlapping regions using t-tests in R.

To test for genetic divergence between SAS and wild fish we first used BisSNP v1.0.0 (Liu *et al.* 2012) to call single nucleotide polymorphisms from the aligned bisulfite sequencing reads. For this analysis we retained all 16 adult samples and one individual per full-sib family from the juvenile dataset (N = 8). Juvenile individuals were chosen to maximize the number of successfully called genotypes. We required a minimum depth of coverage of 8X to call an individual genotype and we retained only SNPs with a successful genotyping rate of 80% (N = 24 individuals). We further removed SNPs with minimum allele frequency of 5% (minimum of 2 alternate alleles). We used AMOVA implemented in pegas v0.13 (Paradis 2010) to test for genome-wide differentiation. We used BayeScan v2.1 (Foll and Gaggiotti 2008) with a liberal prior odds setting of 10 as well as OutFLANK v0.2 (Whitlock and Lotterhos 2015) to identify potential outliers between SAS and wild groups. For the OutFLANK analysis we first built the genome-wide null distribution of divergence based on a linkage pruned set of 12,221 SNPs (obtained with ‘-indep 50 5 1’ using PLINK v0.19) and tested for outliers in the whole dataset based on this distribution. We also employed a polygenic framework to test for subtle correlated changes across many alleles using redundancy analysis (Forester *et al.* 2016) implemented in the vegan v2.5-6 R package (Oksanen *et al.* 2019).

## Data availability

The processed methylation datasets are available in NCBI’s Gene Expression Omnibus (GEO Series accession: GSE162129) and the raw pool-seq and bisulfite-seq data are available at NCBI (Pool-seq: PRJNA679718; bisulfite-seq: PRJNA680707). The ICSASG_v2 Atlantic Salmon genome is available from NCBI (RefSeq accession: GCF_000233375.1). Code to process the data and generate the results are available on github as follows: generation of methylation counts from sequencing data (https://github.com/kylewellband/bwa-meth_pipeline.git), differential methylation analyses (https://github.com/kylewellband/methylUtil.git), C/T polymorphism characterization (https://github.com/kylewellband/CT-poly-wgbs.git). Phenotypic data for juveniles are included in the Supplemental File S1.

## Results

To identify the potential for intergenerational transmission of environmentally-induced variation in DNA methylation acquired during gamete maturation, we used whole-genome bisulfite sequencing to characterize genome-wide DNA methylation variation in six adult smolt-to-adult supplementation (SAS) and six adult natural-origin (wild) Atlantic Salmon (*Salmo salar*) from a smolt-to-adult supplementation program in the Miramichi River in New Brunswick, Canada (Figure 1). Here, SAS individuals originated from a collection of wild juveniles (predominantly age 2 and 3) captured during their migration to salt-water (Figure 1A-B) and reared for two years in land-based freshwater tanks (Figure 1C). Because the zebrafish methylome is reprogrammed to the paternal state (Skvortsova *et al.* 2019), we first compared DNA methylation in sperm cells from maturing male SAS fish to wild male salmon from overlapping cohorts that had spent one year (i.e. grilse) to two years (i.e. multi-sea-winter salmon) in the marine environment (Figure 1D) and were returning to reproduce in freshwater (Figure 1E) to detect the potential for paternally-derived methylation differences as a result of maturation environment. We then created pure-type crosses of SAS and wild adults and reared the offspring in a common environment. We characterized growth-related phenotypic differences in 10-month old juveniles as well as DNA methylation patterns in their livers, an organ known for its important role in regulating growth and metabolism (Trefts *et al.* 2017). Liver also represents a tissue in which a very large number of genes are being expressed in salmonids that could be regulated by epigenetic mechanisms (e.g. Sutherland *et al.* 2019; Rougeux *et al.* 2019) as well as its relative homogeneity of cell-types that could confound methylation analyses (Jaffe and Irizarry 2014), to determine the presence of inherited DNA methylation patterns and their potential influence on proxies of fitness.

### Differential methylation in sperm

For adults we quantified cytosine DNA methylation in sperm for over 16.4 million sites in the cytosine-phosphate-guanine (CpG) context with greater than 5X coverage, but less than 20X coverage in 75% (9 / 12) of samples. Mean coverage per individual was 8.3X (range 4.6X – 19X; Table S1). CpGs in salmon sperm were highly methylated (average methylation: 89%) and exhibited a bimodal distribution, with ~5% of CpGs nearly devoid of methylation (<5%) and 91% of CpGs having methylation levels >80% (Figure S1), consistent with levels reported in other salmonids (Gavery *et al.* 2018) and fish in general (Jiang *et al.* 2013; Potok *et al.* 2013; Ortega-Recalde *et al.* 2019; Skvortsova *et al.* 2019).

We identified differential methylation for individual CpGs between wild and SAS adults using beta-binomial models (Feng *et al.* 2014). There were 4,998 differentially methylated cytosines (DMC; p-value < 0.001) identified between adult SAS and wild salmon sperm that were grouped into 284 differentially methylated regions (DMR; Figure 2A). Regions ranged in size from 51 to 2229 bp, contained between four and 34 CpGs, and comprised 90.2% of DMCs with false discovery rate (FDR) < 0.05. The magnitude of methylation difference between SAS and wild salmon for the identified DMRs averaged 39% (range: 8% to 70%). DMRs in SAS fish were twice as frequently hypo-methylated relative to wild fish than hyper-methylated and this difference was highly significant (68%: 193/284 hypo-DMRs; 32%: 91/284 hyper-DMRs; binomial p-value < 0.001). DMRs overlapped 237 genes or their cis-regulatory contexts (within 5,000 bp of gene features; Table S2). Gene ontology enrichment analysis revealed DMRs in sperm were associated with genes significantly enriched (p-value < 0.05) for a variety of functions in signal transduction pathways, brain development, tissue differentiation, muscle development and contraction, and chromatin silencing (Table S3).

**Figure 2:**
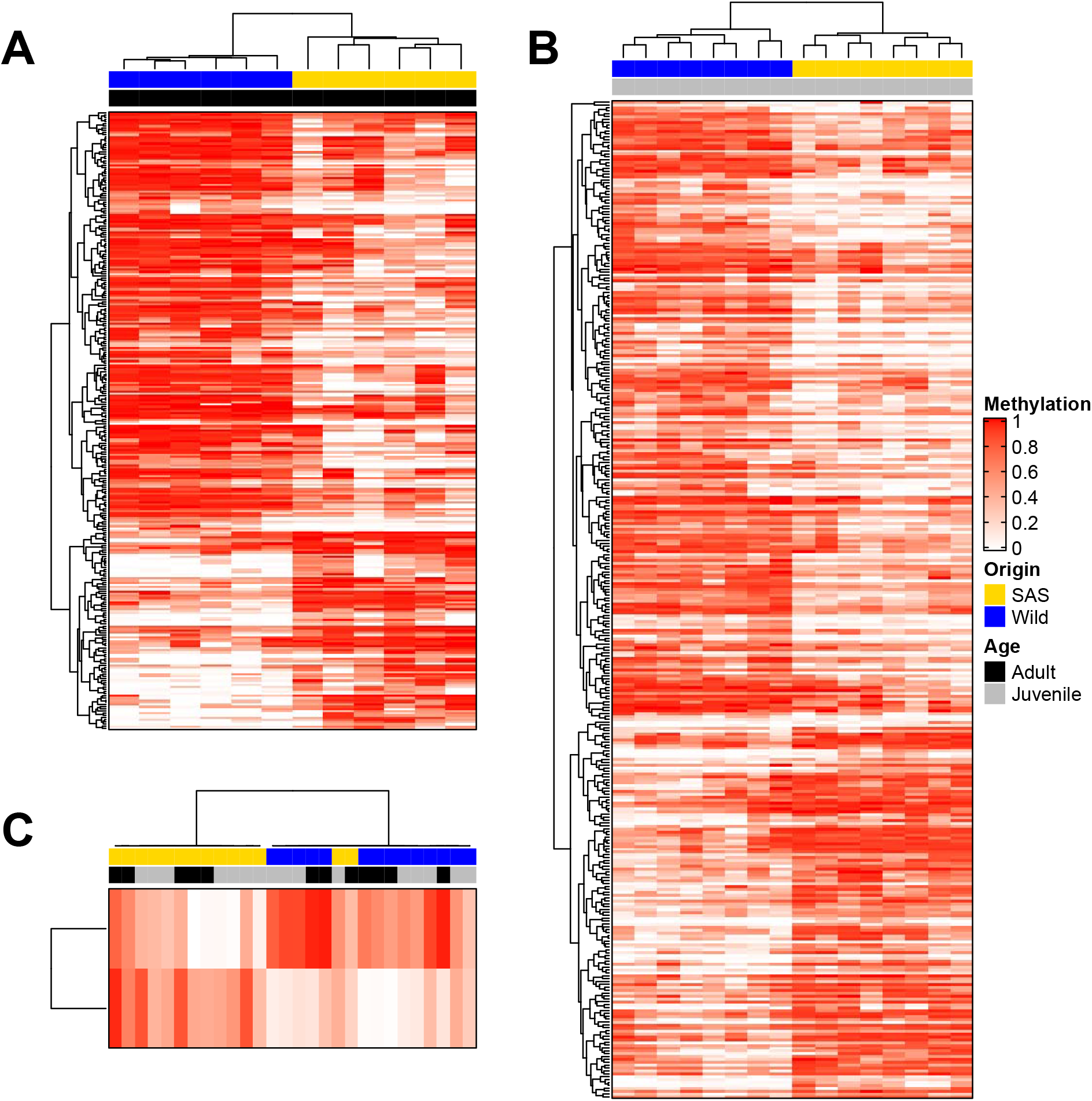
Differentially methylated regions (DMRs; rows) between smolt-to-adult supplementation (SAS; yellow) and wild (blue) salmon. Methylation percentage for each region in each individual (cells of the heatmaps) is expressed as a fraction where un-methylated = 0 (white) to completely methylated = 1 (red). In adult sperm tissues, (A) 284 DMRs were identified (193 hypo-methylated in SAS and 91 hyper-methylated in SAS). In juvenile liver tissues, (B) 346 DMRs were identified (215 hypo-methylated in SAS and 131 hyper-methylated in SAS). Two DMRs (C) were found in common between the two tissues and exhibited similar patterns of differential methylation in both adults and juveniles.

### Differential methylation in juvenile liver

F1 SAS juveniles tended to be longer (SAS: 65.2 ± 6.6 mm, wild: 63.2 ± 6.6 mm; mean ± SD; Figure 3A) and heavier (SAS: 3.42 ± 1.0 g, wild: 3.14 ± 1.0 g; mean ± SD; Figure 3B) than F1 wild juveniles at 10-months of age, but these differences were dwarfed by among-family variation and are not statistically significant when controlling for the number of families investigated (length: F_1,8_ = 0.19, p = 0.68; weight: F_1,8_ = 0.13, p = 0.72). While F1 wild fish were smaller on average, they had similar condition factors (SAS: 1.20 ± 0.06, wild: 1.21 ± 0.06; mean ± SD; F_1,8_ = 2.43, p = 0.16; Figure 3C). Statistically significant patterns consistent with our results have been observed for larger cohorts of Atlantic Salmon in Norway (Skaala *et al.* 2019).

**Figure 3:**
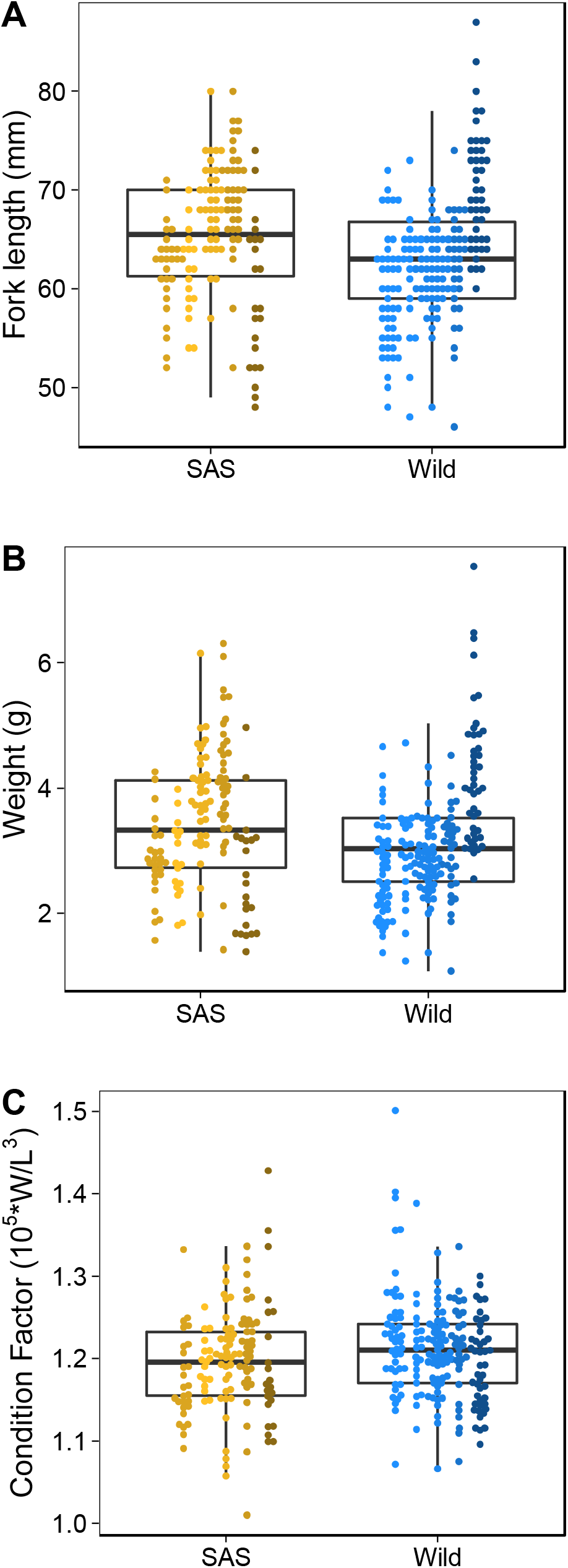
Phenotypic variation of juvenile (10 month old) offspring from five families of smolt-to-adult supplementation (SAS; yellows; N = 138; within-family N: 17 – 37) and five families of wild (blues; N = 198; within-family N: 21 – 53) salmon reared in a common environment. While SAS juveniles were on average longer (A; SAS: 65.2 ± 6.6 mm, Wild: 63.2 ± 6.6 mm) and heavier (B; SAS: 3.42 ± 1.0 g, Wild: 3.14 ± 1.0 g) than wild juveniles when reared in a common hatchery environment, wild juveniles had similar condition (C; SAS: 1.20 ± 0.06, Wild: 1.21 ± 0.06). All measurements are mean ± standard deviation. Controlling for among-family variance using a linear model with a randomeffect for family rendered none of the comparisons statistically significant (length: F_1,8_ = 0.19, p = 0.68; weight: F_1,8_ = 0.13, p = 0.72; condition factor: F_1,8_ = 2.43, p = 0.16).

To investigate the potential for inherited methylation patterns in juvenile liver tissues, we quantified DNA methylation at over 23.1 million CpG sites covered with a minimum of 5X and a maximum of 20X in 75% (12/16) of F1 wild and SAS juveniles. Mean coverage per individual was 9.4X (range: 3.5X to 13.2X; Table S1). We chose two individuals from each of four pure-type families for each treatment group (N = 16). In contrast to sperm, the more metabolically active liver tissues exhibited average methylation levels of approximately 80%. Juvenile liver tissue also exhibited a bimodal distribution of methylation where ~5% of CpGs were unmethylated (<5%), 76% of CpGs with methylation >80%, and a larger fraction of sites with intermediate methylation levels (19% liver CpGs vs. 4% sperm CpGs with methylation fraction between 5 – 80%; Figure S2).

Differentially methylated CpGs (p-value < 0.001; N = 5654) between wild and SAS juvenile offspring were organized into 346 DMRs that ranged in size from 51 to 2131 bp, contained between 4 to 40 CpGs, and covered 98% of DMCs with FDR < 0.05 (Figure 2B). Similar to sperm cells, hypo-methylation was almost twice as common in SAS juvenile liver tissues compared with hyper-methylation and this difference was highly significant (62%: 215/346 hypo-DMRs; 38%: 131/346 hyper-DMRs; binomial p-value < 0.001) and the average magnitude of methylation differences between SAS and wild offspring were comparable (mean: 30%; range: 5 – 52%). DMRs in juvenile liver tissues overlapped 274 genes or their cis-regulatory context. Over-represented biological functions of these genes reflected nervous system development and regulation, muscle development and contraction, signal transduction pathways, and immune system processes (Table S5).

We found overlap of DMRs between the two tissues and life stages (2/622 total DMRs; Figure 2C) that was more than would have been expected by chance (1000 permutations; p < 0.001). The shared regions exhibited the same direction of differential methylation in both tissues (Figure S3A-B) and hierarchical clustering of these regions by individual largely recapitulated the SAS vs. wild groupings (Figure 2C). These regions are in proximity (<20 kb) to genes involved in immune response, tissue differentiation and organ development, and G protein-coupled receptor signaling (Figure S3A-B). Five additional genes were overlapped by DMRs in both tissues but the DMRs in each tissue were not located at the same sites. Of these, metabotropic glutamate receptor 4 (*GRM4*) had regions that were in close proximity (<5,000 bp). The DMRs in proximity to *GRM4* were hypo-methylated in adult SAS sperm and hypermethylated DMR in juvenile SAS liver tissue (Figure S3C).

### Co-methylation network analysis

As a means to investigate whether methylation influences juvenile phenotypic variation (i.e. size and condition factor), we used a network-based approach (Langfelder and Horvath 2008) to identify modules of correlated methylation signatures across the juvenile liver tissue samples and tested for module associations with juvenile phenotypes. To construct the network, we first binned methylation in non-overlapping 100 bp windows and selected the windows (i.e. regions) with among-individual variances greater than 0.05 (N = 59,803 regions). Approximately scale-free networks of correlated methylation regions were constructed using the approach implemented in the R package WGCNA (Langfelder and Horvath 2008). We identified 124 modules that included a total of 4,179 genomic regions. Eighteen modules exhibited significant correlations with at least one phenotype (p < 0.05; Figure S4). Modules were enriched for a variety of signalling pathways and developmental processes relevant to the correlated phenotypes (e.g. growth factor signalling, skeletal muscle development; Table S6). The strongest association existed between the ‘navajowhite1’ module and condition factor (r = 0.69, p = 0.003). The regions in this module were in proximity (5 kb) to genes enriched for mesoderm development and regulation of transcription (Tables S5 and S6). In particular, the gene encoding insulin-like growth factor I (*IGF-1*), a hormone produced in the liver and a key regulator of growth in muscle and skeletal tissues (Ohlsson *et al.* 2009), was associated with this module. Elevated methylation of *IGF-1* in juvenile livers, with an expected reduction in *IGF-1* expression, was associated with reduced weight and length, but better condition in juvenile salmon (Figure S4).

Three modules exhibited significant overlap with DMRs identified between juvenile SAS and wild salmon (permutation tests; p < 0.001). DMR-overlapping regions in these three modules had consistently higher module centrality than non-DMR-overlapping regions (Figure 4; purple: t35.0 = 2.9, p = 0.007; yellow4: t_26.0_ = 3.0, p = 0.006; darkorange: t_24.4_ = 2.6, p = 0.01). Overall, these results indicate that 1) juvenile phenotypes are influenced by methylation variation among individuals and, 2) hatchery-induced differential methylation affects key loci that are central to certain modules influencing offspring phenotypes.

**Figure 4:**
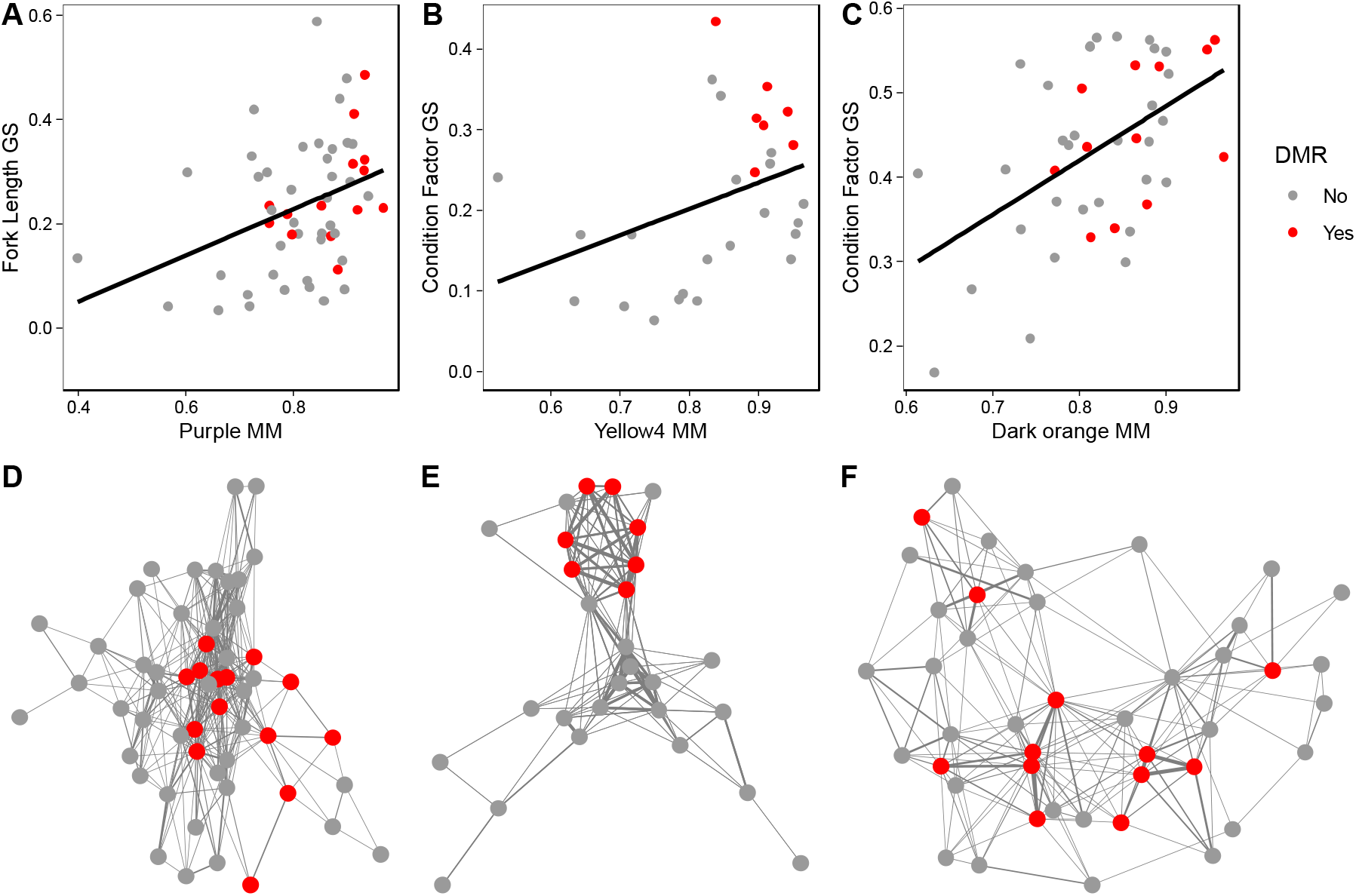
Differentially methylated regions (DMR) between SAS and wild salmon that overlap juvenile phenotype-associated methylation module regions (circles: red = DMR overlap, grey = no overlap) are over represented in module cores. Three methylation modules correlated with juvenile phenotypes (A & D: purple – weight, r = −0.34, p = 0.04; B & E: yellow4 – condition factor, r = −0.23, p = 0.04; C & F: darkorange – condition factor, r = −0.53, p = 0.03). All modules exhibited a significant correlation between module membership (x-axes; MM = absolute value of the correlation of region methylation with the main axis of module variation) and gene significance (y-axes; GS = absolute value of the correlation of region methylation with the phenotype). DMR overlapping regions in these modules were more highly correlated with the main axis of variation of the modules than non-DMR overlapping regions (A – purple: t_35.0_ = 2.9, p = 0.007; B – yellow4: t_26.0_ = 3.0, p = 0.006; C – darkorange: t_24.4_ = 2.6, p = 0.01). DMR overlapping regions were also more centrally located and highly connected in module networks than non-DMR overlapping regions (D – F).

To account for the possibility of selection causing genetic differences between juvenile SAS and wild salmon and explaining the observed phenotypic differences, we also genotyped individuals for 974,219 single nucleotide polymorphisms (excluding CpG context C/T and A/G SNPs to avoid confounding methylation variation with allelic variation) using the aligned bisulfite sequencing reads and performed outlier tests. Wild salmon may experience selection during their time in the marine environment (i.e. smolt to adult; Bourret *et al.* 2014) where mortality can range from 65 to 99% (Chaput 2012). In contrast, SAS salmon may experience relaxed selection during this time in the hatchery with a mortality rate from smolt to adult of approximately 40%. Overall, we failed to find support for a genome-wide average FST larger than zero (AMOVA: 1000 permutations, p = 0.77). Using outlier detection algorithms, we did not detect significant shifts in allele frequencies between SAS and wild salmon using BayeScan or a polygenic framework (RDA; R^2^ = 0, p = 0.71), and we only identified two outlier SNPs with FDR < 0.01 in OutFLANK (Figure S5; Table S8), suggesting an overall absence of differential selection at the genome level between SAS and wild salmon.

## Discussion

### Maturation environment influences gamete methylation

Our results demonstrate that environmental variation (i.e. growth and maturation in a natural vs. hatchery setting) experienced after approximately two years of common rearing in their natural riverine environment alters both DNA methylation in salmon sperm as well as DNA methylation of hatchery-produced progeny with SAS parents. The maturation-environment effect we demonstrated strongly suggests, as other have reported (e.g. Dias and Ressler 2014; Sciamanna *et al.* 2019), that the Weismann Barrier is permeable and that information perceived by the soma can be incorporated into the germline via epigenetic mechanisms. To our knowledge however, this is the first-time environmental variation experienced later in life has been directly demonstrated to influence gamete DNA methylation in fish.

Several lines of evidence support a mechanism of environmentally-mediated DNA methylation remodelling during gamete maturation. First, environmental differences in early life are known to influence gamete methylation (Gavery *et al.* 2018; Rodriguez Barreto *et al.* 2019). Second, multiple copies of DNA methyltransferase 3 (*DMNT3*) the methyltransferase responsible for the addition of new DNA methylation, have been retained following successive genome-duplication events (i.e. ohnologs) for salmonid and other teleost fishes (Liu *et al.* 2020). In Rainbow Trout (*Oncorhynchus mykiss*), certain *DMNT3* ohnologs are expressed in spermatozoa during late spermatogenesis (i.e. a few weeks before spawning; Liu *et al.* 2020), thus providing a mechanism by which salmonids may alter DNA methylation in their gametes until days or weeks before spawning. Third, teleost fishes do not appear to experience genomewide reprogramming of paternal methylation patterns following fertilization (Jiang *et al.* 2013; Potok *et al.* 2013) or differentiation of gonadal tissue (Ortega-Recalde *et al.* 2019; Skvortsova *et al.* 2019). While recent evidence from another fish species (*Oryzias latipes*) indicates it does undergo global methylation reprogramming (Wang and Bhandari 2019), the presence of shared differentially methylated regions between our adult and juvenile datasets suggests that at least some genomic regions escape reprograming if it does also occur in salmon. Thus, adult salmon may be able to transmit heritable information to their offspring about the physical or biological environments they experience immediately prior to spawning. As such, our results are consistent with the hypothesis that epigenetic mechanisms can facilitate transgenerational plasticity (Bell and Hellmann 2019).

Transgenerational plasticity is theorized to evolve when environmental variability is sufficiently stable or predictable such that adults can transmit relevant information about the environment to their offspring (McNamara *et al.* 2016). If only early-life epigenetic signals were capable of being transmitted intergenerationally, transgenerational plasticity may not have been expected to evolve for salmon whose lives generally span both temporally and spatially diverse environments, and whose life histories involve limited parental care (Thorstad *et al.* 2010). Our demonstration of the potential for adults to transmit environmental information acquired later in life to their offspring suggests transgenerational plasticity in salmon may be an important factor contributing to life history variation and adaptive responses to environmental change.

### Origins of hatchery-induced differential methylation

There is an unresolved question of whether epigenetic differences arising as a result of hatchery exposure occur due to deterministic processes (i.e. adaptive responses potentially arising from existing molecular machineries as a result of past selection) or stochastic processes (i.e. random environmental perturbations of wildtype methylation patterns). Several lines of evidence suggest the patterns we observed originate at least partly from deterministic processes. First, if methylation changes were completely stochastic we would expect no bias in the direction of methylation changes. In contrast, our results show that differential methylation in SAS fish is strongly biased (about two-fold) toward hypomethylation in both sperm and juvenile livers. Second, despite the fact that our juveniles and adults originated from different cohorts and that juvenile livers will have undergone tissue-specific methylation reprogramming from the state observed in sperm, we detected more DMRs in common between the datasets than expected by chance. Finally, the regions in phenotypically correlated methylation networks that overlap SAS vs. wild DMRs are significantly biased toward being centrally located regions in the comethylated networks indicating these are not random associations.

More generally, prior research has established parallel signatures of DNA methylation divergence in response to early-life hatchery rearing in two populations of Coho Salmon (*Oncorhynchus kisutch*; Le Luyer *et al.* 2017) and hatchery-reared Rainbow Trout exhibit a significant proportion (approximately 20%) of hatchery-origin DMRs that are shared between red blood cells and sperm (Gavery *et al.* 2018). There are similarities (discussed in detail below) in the biological functions impacted by the epigenetic signatures of response to early-life hatchery rearing for Rainbow Trout (Gavery *et al.* 2018), Coho Salmon (Le Luyer *et al.* 2017), and Atlantic Salmon (Rodriguez Barreto *et al.* 2019). These epigenetic signatures of early-life hatchery rearing are also broadly similar to epigenetic signatures of domestication observed in recently domesticated European Sea Bass (*Dicentrarchus labrax*; Anastasiadi and Piferrer 2019).

### Epigenetic signatures of domestication

We identified seven genes associated with SAS versus wild differential methylation that have been reported in previous studies of hatchery induced differential methylation in salmonids. Phosphatidylinositol 3-kinase regulatory subunit alpha (*PIK3R1*) was differentially methylated in SAS sperm cells and has previously been identified as differentially methylated in sperm of Atlantic Salmon reared in a hatchery from birth (Rodriguez Barreto *et al.* 2019). This gene is an important regulator at the center of many growth factor and hormone signalling pathways (Cantley 2002) and has also previously been identified as potentially target of selection in domesticated salmon (Liu *et al.* 2017). Three additional genes reported in previous studies that we report as differentially methylated in sperm cells all have roles in the nervous system including growth and differentiation (*NRG2*; Britsch 2007), cell-cell adhesion (*PCDHGC5*; Wang *et al.* 2002), and neurotransmitter release (*STXBP5L*; Geerts *et al.* 2015).

The remaining three genes we detected in common with other studies were all differentially methylated in juvenile liver tissue with reported functions in other organisms of epidermal tissue development (*BCR*; Dubash *et al.* 2013), cell adhesion and cytoskeleton organization during neuron differentiation (*CTNNA2*; Schaffer *et al.* 2018), and neuron differentiation (*ARHGAP32*; Nakamura *et al.* 2002). *BCR* has previously been identified as differentially methylated in hatchery-origin Coho Salmon white muscle (Le Luyer *et al.* 2017), *CTNNA2* in both red blood cells and sperm of hatchery-origin Rainbow Trout (Gavery *et al.* 2018), and *ARHGAP32* in sperm of Atlantic Salmon (Rodriguez Barreto *et al.* 2019). In general, patterns of DNA methylation across studies implicate regulation of cell differentiation and developmental processes with a particular enrichment of genes involved in neuron differentiation.

We identified several glutamate receptors as being differentially methylated either in sperm (*GRIK5, GRM4*) or liver (*GRM4, GRID2, GRM3*). Glutamate receptors are well known targets of selection in many domestic animals (O’Rourke and Boeckx 2020) and have specifically been identified as being differentially methylated between wild and domestic European Sea Bass (Anastasiadi and Piferrer 2019). Glutamatergic signalling is an important excitory driver of the hypothalamic–pituitary–adrenal (HPA) axis that, among other functions, mediates organismal stress responses and aggression and it has been hypothesized that selection on these pathways underlies “tameness” in domesticated animals and attenuated stress responses under crowded conditions (O’Rourke and Boeckx 2020). This raises the hypothesis that adult SAS fish could transmit information about the crowding they experienced prior to spawning to their offspring in order to prime them for a highly competitive environment upon hatching (Christie *et al.* 2012).

In summary, were the epigenetic effects induced by hatchery environments truly random, it would be very unlikely to detect particular genes or pathways across multiple studies as revealed by the available literature. Altogether, the compiled evidence supports the hypothesis that there is a certain degree of conservation in the DNA methylation changes in response to captive rearing across a broader taxonomy of teleost fishes.

### Consequences of hatchery-induced methylation for offspring phenotypes

We identified several correlated methylation profiles that were statistically associated with offspring phenotypes. Conceptually, this analysis identifies pathways or biological functions that are co-regulated by methylation. Of the 18 methylation modules that were correlated with juvenile phenotypes, several occurred in proximity to genes enriched for signalling pathways (i.e. muscle growth and differentiation, skeletal development, neural development, and immune system processes) directly relevant to the phenotypes being studied. In particular, the methylation module in juvenile livers with the strongest phenotypic correlation (i.e. navajowhite1) contained regions overlapping *IGF-1. IGF-1* is a hormone produced and released from the liver in response to growth hormone (*GH*) signalling that plays a key endocrine role mediating growth and differentiation of muscle and skeletal tissues and therefore body size (Ohlsson *et al.* 2009). Body size and condition are traits closely linked with juvenile salmonid survival and fitness (Quinn and Peterson 1996; Einum and Fleming 2000). In addition, the *GH/IGF-1* axis has been implicated in acclimation to saltwater (McCormick 2001) which is a major selective barrier for anadromous salmonids and has been identified as a deficiency of hatchery reared fish (Shrimpton *et al.* 1994). Our results demonstrate that differences in the DNA methylation state of this important growth-regulating gene have the potential to exert influence on salmon growth and developmental trajectories that are likely to have real consequences for individual fitness.

Hatchery-induced differential methylation may directly influence both the fitness-related traits we quantified here and other more complex behavioral traits with fitness consequences at later stages of development than we have studied. Hatchery-induced DMRs overlapped regions centrally located in co-methylated modules that were associated with biological functions involved in brain neuron differentiation as well as the glutamate receptor *GRM4* (i.e. module yellow4) suggesting that hatchery-environment-induced methylation-mediated behavioural changes (e.g. attenuated stress response to crowding; O’Rourke and Boeckx 2020) could have consequences for the growth trajectories of offspring. We also identified modules (i.e. purple) and differential methylation of genes not included in methylation modules that lie downstream of genes in important phenotype-associated modules (i.e. *IGF-1* signalling pathway). Phosphatidylinositol-mediated signaling (i.e. module purple) and *PI3KR1* in particular plays a key role in modulating the response to *IGF-1* stimulation (Hakuno and Takahashi 2018) and thus hatchery-induced differential methylation may influence or modulate growth trajectories at a later step in that signalling cascade that, at least in part, may explain the subtle differences we observed in phenotypes between SAS and wild juveniles. Collectively, our results suggest that hatchery-associated effects could indeed be mediated through DNA methylation with consequences for aspects of fish phenotypes with known relationships to their fitness.

### Comparison of hatchery rearing approaches

Despite detecting DNA methylation differences between SAS and wild fish, our work also reveals some fundamental differences between the effects of early-rearing and later-life hatchery exposure for salmon. SAS fish (both SAS adults and their progeny) exhibited more hypo-methylation relative to wild fish (for both adult males and their progeny reared in a common environment), in contrast to previous work that has demonstrated predominantly hypermethylation of fish produced and reared in hatchery from the egg stage (Le Luyer *et al.* 2017; Gavery *et al.* 2018; Rodriguez Barreto *et al.* 2019). Reduced representation bisulfite sequencing (RRBS) in Coho Salmon and Rainbow Trout reported differential methylation at between 0.03% and 0.1% of analyzed sites or regions. Using similar criteria, our results showed differential methylation affected an order of magnitude less CpG sites (0.004%). This suggests that the potential for DNA methylation-mediated domestication effects caused by later-life hatchery exposure may not be as severe as those observed for salmon that experience early-life hatchery rearing. RRBS approaches are believed to preferentially target gene regulatory relevant regions of the genome (e.g. Anastasiadi and Piferrer 2019) and thus, because of a difference in techniques between studies (RRBS vs. whole-genome bisulfite sequencing), our comparisons may be biased. However, the genomic distribution and density of regulatory relevant CpGs in non-mammalian vertebrates fundamentally differs from that of mammals (Long *et al.* 2013). Bioinformatic interrogation of our data indicates RRBS applied to our study would have assayed approximately 7 million CpGs and only detected 10% of the observed DMCs implying the above comparison is reasonably unbiased. Furthermore, it suggests that in salmon, and possibly fishes more generally, RRBS approaches fail to capture a significant proportion of biologically relevant methylation differences.

Like previous studies, we have demonstrated a lack of genome-wide sequence differentiation between hatchery-reared and wild fish (Christie *et al.* 2016; Le Luyer *et al.* 2017; Gavery *et al.* 2018). This result suggests that the selective regime imposed by the hatchery environment over one generation was not strong enough to cause widespread differentiation. In turn, it also suggests that despite the high levels of mortality during the marine phase of salmons’ lives (65 – 99%; Chaput 2012), selection in the marine environment may not be important enough to cause widespread, temporally consistent, changes in allele frequencies between wild and SAS salmon. Previous work in Atlantic Salmon has reported consistent allele frequency changes over the marine migration period for only one of two populations studied indicating patterns of differentiation due to the marine environment are spatially and temporally variable (Bourret *et al.* 2014). It is difficult to know if the two outliers we detected result from selection in the hatchery or marine environments. In spite of this, our results clearly implicate a stronger role for epigenetic factors and not differences in genetic variation in hatchery-related phenotypic divergence.

We have demonstrated the potential for environmental effects to be propagated to offspring for salmon who experience hatchery environments during maturation via the intergeneration transmission of DNA methylation. Our experiments on juvenile fish were conducted in a laboratory setting and thus whether these effects are also detectable and have fitness consequences in the true context of SAS program where SAS individuals are released and reproduce naturally in the wild remains unknown. Genotype-by-environment interactions are pervasive in salmonids (Vandersteen *et al.* 2019) and so are epigenotype-by-environment interactions (Christensen *et al.* 2021). As such, there is an urgent need to evaluate the interaction between SAS and wild rearing on offspring DNA methylation and development in a natural environment. Other sources of epigenetic information (i.e. small RNAs) are also well known to mediate intergenerational effects (Sciamanna *et al.* 2019) that may well mediate the phenotypic differences between individuals we have observed. On the other hand, multiple epigenetic mechanisms often function together to affect phenotypic changes (Cavalli and Heard 2019) and future work unravelling the mechanistic basis of hatchery-induced phenotypic effects will need to clarify the potential contributions of other epigenetic mechanisms, their relative importance, and the degree to which these effects are reversible following the cessation of the environmental exposure. Only then will the evolutionary consequences of environmentally-induced epigenetic variation in these systems be globally understood.

## Acknowledgements

We sincerely thank Mark Hambrook and staff from the Miramichi Salmon Association who provided logistical support for the capture and rearing of juvenile salmon and the collection of wild adults. We also thank three anonymous referees for their constructive comments on a previous version of this manuscript. Financial and logistical support for this work was provided by the Collaboration for Atlantic Salmon Tomorrow Inc. including Cooke Aquaculture Inc., J.D. Irving Ltd. and the Atlantic Canada Opportunities Agency. Funding for this work was also provided by the Canadian Research Chair in Genomics and Conservation of Aquatic Resources, as well as MITACS, and an NSERC Postdoctoral Fellowship to KW.

## Author contributions

L.B., T.L., R.A.C., and K.W. conceived the study. T.L. and R.A.C. secured the funding. K.W. and L.B. designed the study. D.R. coordinated rearing of the fish. D.R. and K.W. collected the samples. K.W. analyzed the data and wrote the first draft. All authors revised and approved of the final version.

## Supplemental information

**Supplemental Table S1:** Statistics of sequencing effort and coverage of whole genome bisulfite sequencing for all samples.

**Supplemental Table S2:** Genes associated with differentially methylated regions in sperm.

**Supplemental Table S3:** Gene ontology biological process terms associated with differentially methylated regions in sperm.

**Supplemental Table S4:** Genes associated with differentially methylated regions in juvenile liver.

**Supplemental Table S5:** Gene ontology biological process terms associated with differentially methylated regions in juvenile liver.

**Supplemental Table S6:** Gene ontology biological process terms associated with methylated regions included in co-methylated modules.

**Supplemental Table S7:** Genes associated with methylated regions included in co-methylated modules.

**Supplemental Table S8:** Genomic location of two outlier SNPs detected at a FDR < 0.05 between SAS and wild fish using OutFLANK.

**Supplemental File S1:** Phenotypic data for lab reared juveniles.

**Supplemental Figure 1:**
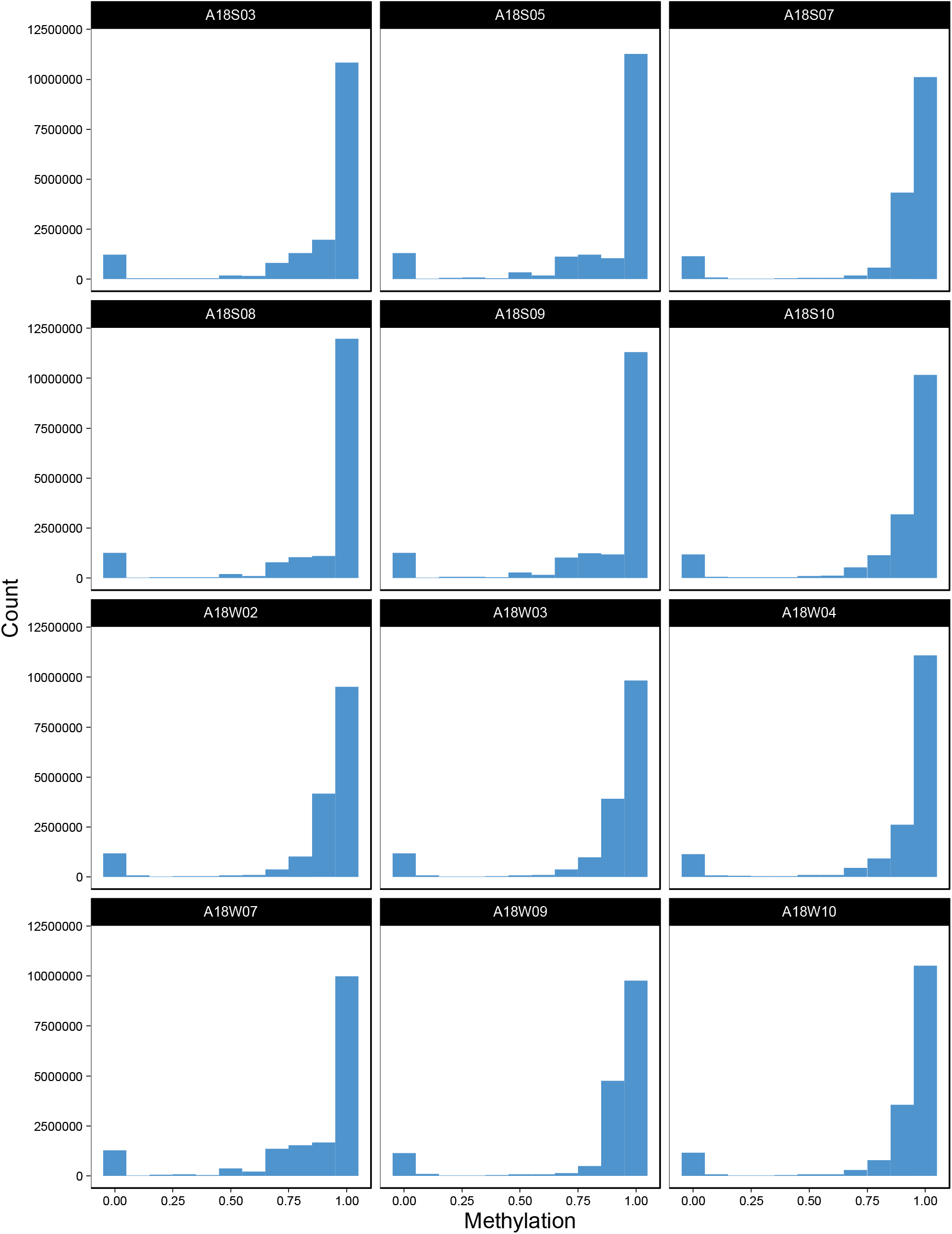
Distribution of methylation proportion (# methylated bases / read coverage) for 16.4 million CpG sites from adult sperm samples.

**Supplemental Figure 2:**
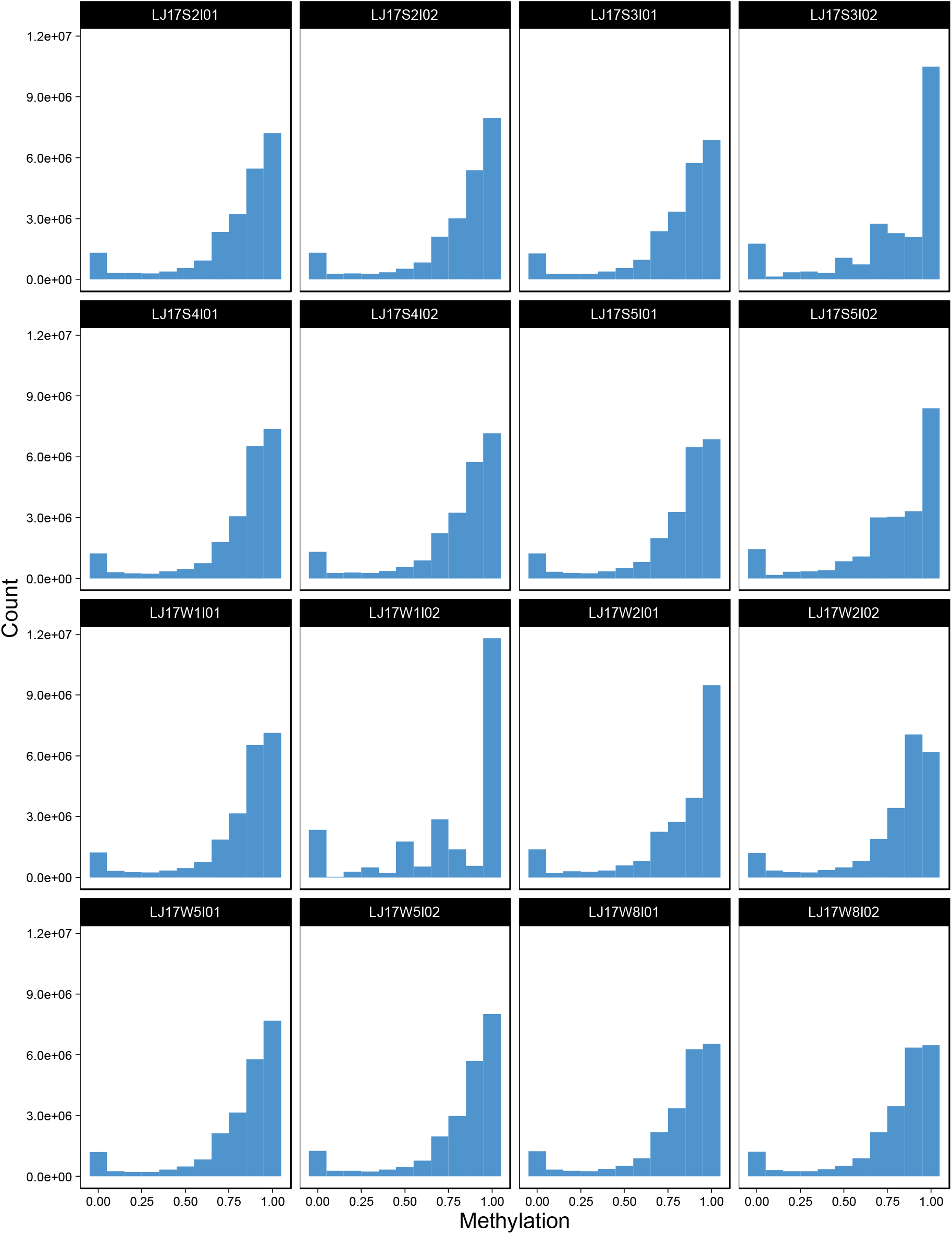
Distribution of methylation proportion (# methylated bases / read coverage) for 23.1 million CpG sites from juvenile liver samples.

**Supplemental Figure 3:**
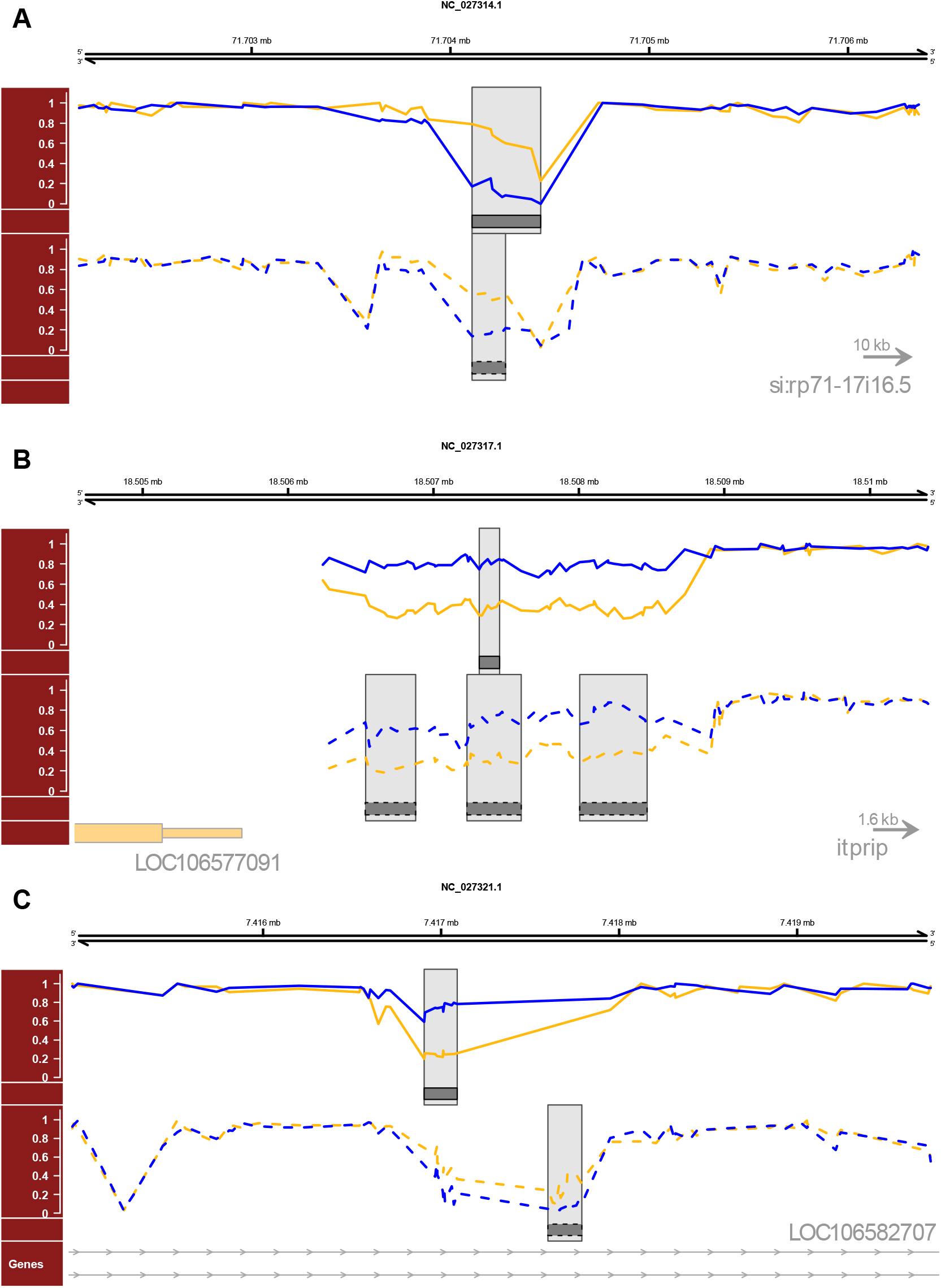
Differential methylated regions (DMR) between SAS (yellow) and wild (blue) salmon that overlapped (A-B) or targeted the same gene (C) between sperm (solid lines) and juvenile liver tissues (dashed lines). Grey boxes highlight the extent of DMRs and the lower tracks indicate annotated genes.

**Supplemental Figure 4:**
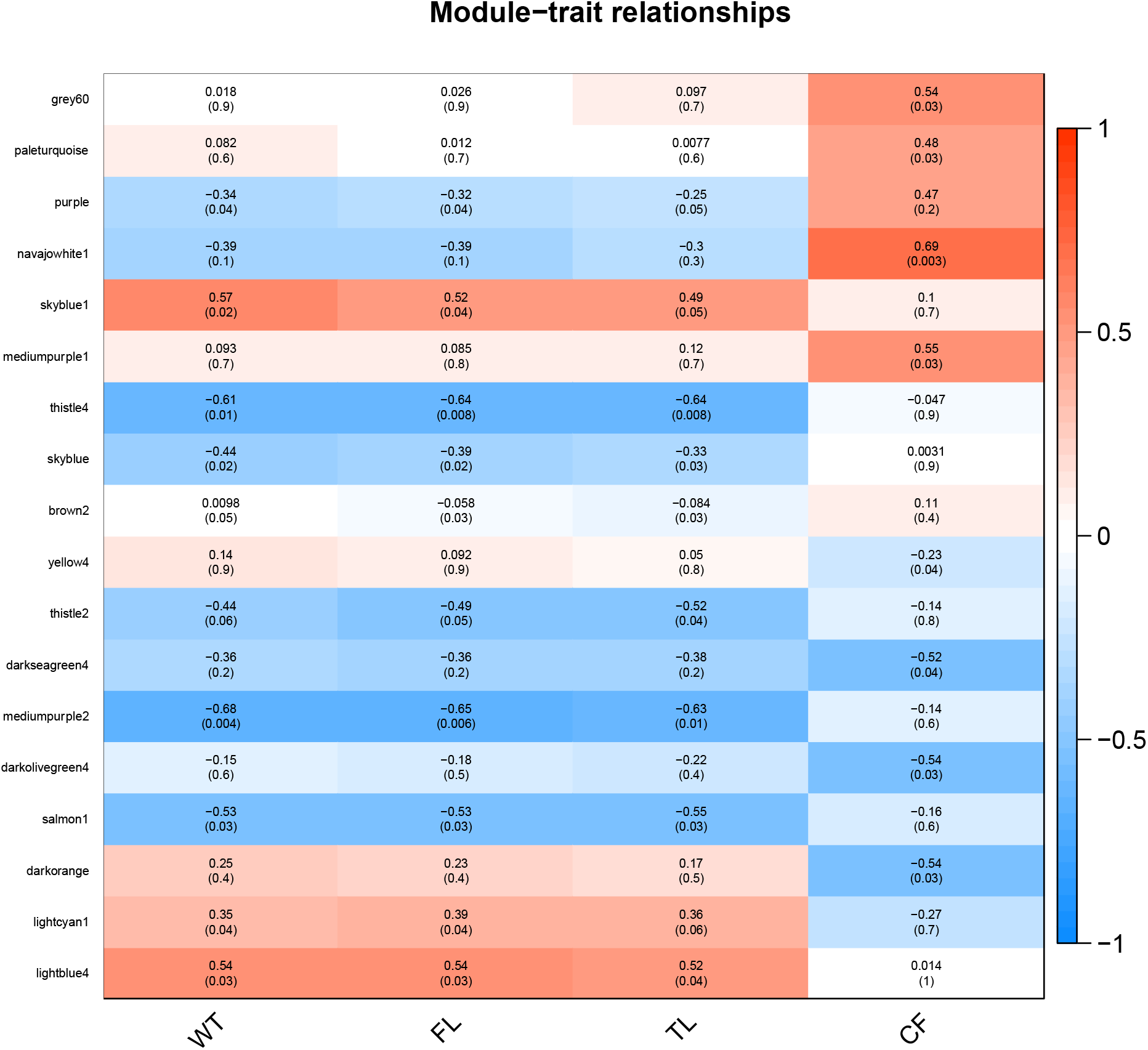
Heatmap of methylation – phenotype correlations for all modules (represented by color names on the y-axis) with at least one significant correlation. Colors and values in the cells represent the magnitude and direction of the correlation coefficient (blue = negative correlation, red = positive correlation) with the statistical significance of the correlation coefficient in parentheses (p-value). Phenotypes on the x-axis: WT = weight in g, FL and TL = fork length and total length in mm, and CF = condition factor (10^5^ * WT / FL^3).

**Supplemental Figure 5:**
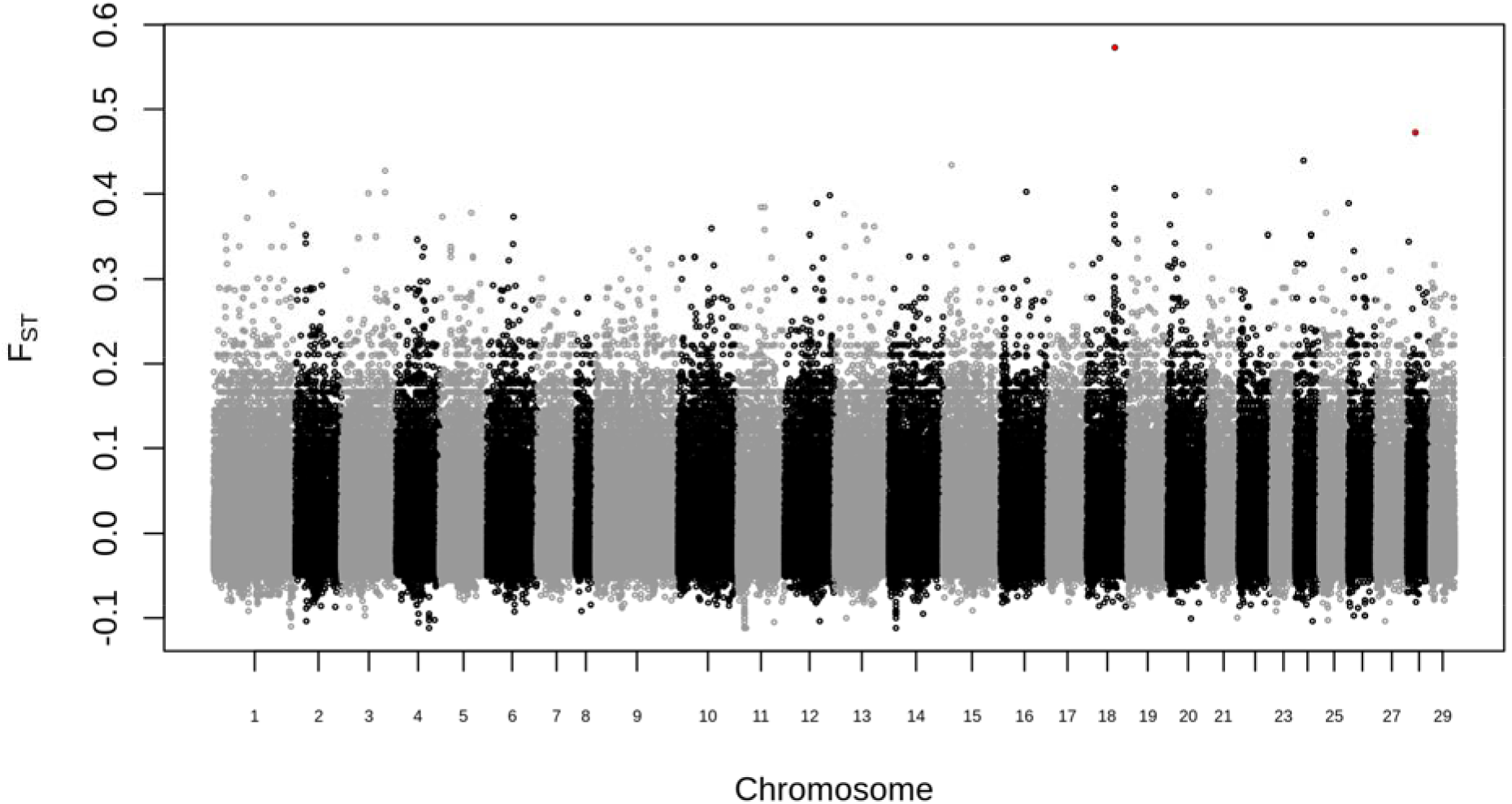
Distribution of genetic divergence (F_ST_) between SAS and wild salmon for 974,219 single nucleotide polymorphisms. The genome-wide average divergence (F_ST_) did not differ from zero (AMOVA; 1000 permutations: p = 0.77). Outliers (FDR < 0.01) identified by OutFLANK are highlighted in red. Neither BayeScan, nor a polygenic framework (RDA, R^2^ = 0, p = 0.71) identified any outliers.

## References

Alexa, A., and J. Rahnenfuhrer, 2019 topGO: Enrichment Analysis for Gene Ontology.

Alexa, A., J. Rahnenführer, and T. Lengauer, 2006 Improved scoring of functional groups from gene expression data by decorrelating GO graph structure. Bioinformatics 22: 1600–1607.

Aljanabi, S. M., and I. Martinez, 1997 Universal and rapid salt-extraction of high quality genomic DNA for PCR-based techniques. Nucleic Acids Res. 25: 4692–4693.

Anastasiadi, D., and F. Piferrer, 2019 Epimutations in Developmental Genes Underlie the Onset of Domestication in Farmed European Sea Bass. Mol. Biol. Evol. 36: 2252–2264.

Araki, H., B. A. Berejikian, M. J. Ford, and M. S. Blouin, 2008 Fitness of hatchery-reared salmonids in the wild. Evol. Appl. 1: 342–355.

Artemov, A. V., N. S. Mugue, S. M. Rastorguev, S. Zhenilo, A. M. Mazur et al., 2017 Genome-Wide DNA Methylation Profiling Reveals Epigenetic Adaptation of Stickleback to Marine and Freshwater Conditions. Mol. Biol. Evol. 34: 2203–2213.

Bell, A. M., and J. Hellmann, 2019 An Integrative Framework for Understanding the Mechanisms and Multigenerational Consequences of Transgenerational Plasticity. Annu. Rev. Ecol. Evol. Syst. 50: 1–22.

Benjamini, Y., and Y. Hochberg, 1995 Controlling the False Discovery Rate: A Practical and Powerful Approach to Multiple Testing. J. R. Stat. Soc. B 57: 289–300.

Berbel-Filho, W. M., N. Berry, D. Rodríguez-Barreto, S. Rodrigues Teixeira, C. Garcia de Leaniz et al., 2020 Environmental enrichment induces intergenerational behavioural and epigenetic effects on fish. Mol. Ecol. 29: 2288–2299.

Berejikian, B. A., and D. M. Van Doornik, 2018 Increased natural reproduction and genetic diversity one generation after cessation of a steelhead trout (*Oncorhynchus mykiss*) conservation hatchery program (H. Wang, Ed.). PLoS One 13: e0190799.

Berejikian, B., D. Van Doornik, A. LaRae, S. Tezak, and J. Lee, 2005 The Effects of Exercise on Behavior and Reproductive Success of Captively Reared Steelhead. Trans. Am. Fish. Soc. 134:1236–1252.

Berejikian, B. A., E. P. Tezak, and S. L. Schroder, 2001 Reproductive Behavior and Breeding Success of Captively Reared Chinook Salmon. North Am. J. Fish. Manag. 21: 255–260.

Bourret, V., M. Dionne, and L. Bernatchez, 2014 Detecting genotypic changes associated with selective mortality at sea in Atlantic salmon: polygenic multilocus analysis surpasses genome scan. Mol. Ecol. 23: 4444–4457.

Britsch, S., 2007 The neuregulin-I/ErbB signaling system in development and disease. Adv. Anat. Embryol. Cell Biol. 190: 1–65.

Broad Institute, 2019 Picard Toolkit. GitHub Repos.

Cantley, L. C., 2002 The Phosphoinositide 3-Kinase Pathway. Science (80-.). 296: 1655–1657.

Cavalli, G., and E. Heard, 2019 Advances in epigenetics link genetics to the environment and disease. Nature 571: 489–499.

Chaput, G., 2012 Overview of the status of Atlantic salmon (*Salmo salar*) in the North Atlantic and trends in marine mortality. ICES J. Mar. Sci. 69: 1538–1548.

Chen, S., Y. Zhou, Y. Chen, and J. Gu, 2018 fastp: an ultra-fast all-in-one FASTQ preprocessor. Bioinformatics 34: i884–i890.

Christensen, K. A., J. Le Luyer, M. T. T. Chan, E. B. Rondeau, B. F. Koop et al., 2021 Assessing the effects of genotype-by-environment interaction on epigenetic, transcriptomic, and phenotypic response in a Pacific salmon (B. J. Andrews, Ed.). G3 Genes|Genomes|Genetics 11:.

Christie, M. R., M. J. Ford, and M. S. Blouin, 2014 On the reproductive success of early-generation hatchery fish in the wild. Evol. Appl. 7: 883–896.

Christie, M. R., M. L. Marine, S. E. Fox, R. A. French, and M. S. Blouin, 2016 A single generation of domestication heritably alters the expression of hundreds of genes. Nat. Commun. 7: 1–6.

Christie, M. R., M. L. Marine, R. A. French, and M. S. Blouin, 2012 Genetic adaptation to captivity can occur in a single generation. Proc. Natl. Acad. Sci. 109: 238–242.

Dias, B. G., and K. J. Ressler, 2014 Parental olfactory experience influences behavior and neural structure in subsequent generations. Nat. Neurosci. 17: 89–96.

Dubash, A. D., J. L. Koetsier, E. V. Amargo, N. A. Najor, R. M. Harmon et al., 2013 The GEF Bcr activates RhoA/MAL signaling to promote keratinocyte differentiation via desmoglein-1. J. Cell Biol. 202: 653–666.

Duempelmann, L., M. Skribbe, and M. Bühler, 2020 Small RNAs in the Transgenerational Inheritance of Epigenetic Information. Trends Genet. 36: 203–214.

Eaton, S. A., N. Jayasooriah, M. E. Buckland, D. I. K. Martin, J. E. Cropley et al., 2015 Roll over Weismann: Extracellular vesicles in the transgenerational transmission of environmental effects. Epigenomics 7: 1165–1171.

Einum, S., and I. A. Fleming, 2000 Selection against late emergence and small offspring in Atlantic Salmon (*Salmo salar*). Evolution (N. Y). 54: 628–639.

Feng, H., K. N. Conneely, and H. Wu, 2014 A Bayesian hierarchical model to detect differentially methylated loci from single nucleotide resolution sequencing data. Nucleic Acids Res. 42: e69–e69.

Foll, M., and O. Gaggiotti, 2008 A Genome-Scan Method to Identify Selected Loci Appropriate for Both Dominant and Codominant Markers: A Bayesian Perspective. Genetics 180: 977–993.

Forester, B. R., M. R. Jones, S. Joost, E. L. Landguth, and J. R. Lasky, 2016 Detecting spatial genetic signatures of local adaptation in heterogeneous landscapes. Mol. Ecol. 25: 104–120.

Fraser, D. J., 2016 Risks and benefits of mitigating low marine survival in wild salmon using smolt-to-adult captive-reared supplementation. Can. Sci. Advis. Secr. Res. Doc. 030: 1–31.

Garrison, E., and G. Marth, 2012 Haplotype-based variant detection from short-read sequencing. arXiv 1207.3907.

Gavery, M. R., K. M. Nichols, B. A. Berejikian, C. P. Tatara, G. W. Goetz et al., 2019 Temporal dynamics of DNA methylation patterns in response to rearing juvenile steelhead (*Oncorhynchus mykiss*) in a hatchery versus simulated stream environment. Genes (Basel). 10:.

Gavery, M. R., K. M. Nichols, G. W. Goetz, M. A. Middleton, and P. Swanson, 2018 Characterization of Genetic and Epigenetic Variation in Sperm and Red Blood Cells from Adult Hatchery and Natural-Origin Steelhead, *Oncorhynchus mykiss*. G3&#58; Genes|Genomes|Genetics 8: g3.200458.2018.

Geerts, C. J., J. J. Plomp, B. Koopmans, M. Loos, E. M. van der Pijl et al., 2015 Tomosyn-2 is required for normal motor performance in mice and sustains neurotransmission at motor endplates. Brain Struct. Funct. 220: 1971–1982.

Hakuno, F., and S.-I. Takahashi, 2018 40 YEARS OF IGF1: IGF1 receptor signaling pathways. J. Mol. Endocrinol. 61: T69–T86.

Heard, E., and R. A. Martienssen, 2014 Transgenerational epigenetic inheritance: Myths and mechanisms. Cell 157: 95–109.

Heckwolf, M. J., B. S. Meyer, R. Häsler, M. P. Höppner, C. Eizaguirre et al., 2020 Two different epigenetic information channels in wild three-spined sticklebacks are involved in salinity adaptation. Sci. Adv. 6:.

Horsthemke, B., 2018 A critical view on transgenerational epigenetic inheritance in humans. Nat. Commun. 9: 1–4.

Jaffe, A. E., and R. A. Irizarry, 2014 Accounting for cellular heterogeneity is critical in epigenome-wide association studies. Genome Biol. 15: R31.

Jiang, L., J. Zhang, J.-J. Wang, L. Wang, L. Zhang et al., 2013 Sperm, but Not Oocyte, DNA Methylome Is Inherited by Zebrafish Early Embryos. Cell 153: 773–784.

Jones, O. R., and J. Wang, 2010 COLONY: a program for parentage and sibship inference from multilocus genotype data. Mol. Ecol. Resour. 10: 551–555.

Jun, G., M. K. Wing, G. R. Abecasis, and H. M. Kang, 2015 An efficient and scalable analysis framework for variant extraction and refinement from population-scale DNA sequence data. Genome Res. 25: 918–925.

Karimzadeh, M., C. Ernst, A. Kundaje, and M. M. Hoffman, 2018 Umap and Bismap: quantifying genome and methylome mappability. Nucleic Acids Res. 46: e120.

Langfelder, P., and S. Horvath, 2008 WGCNA: an R package for weighted correlation network analysis. BMC Bioinformatics 9: 559.

Leitwein, M., M. Laporte, J. Le Luyer, K. Mohns, E. Normandeau et al., 2021 Epigenomic modifications induced by hatchery rearing persist in germ line cells of adult salmon after their oceanic migration. Evol. Appl. eva.13235.

Li, H., 2013 Aligning sequence reads, clone sequences and assembly contigs with BWA-MEM. arXiv Prepr. arXiv00:3.

Lien, S., B. F. Koop, S. R. Sandve, J. R. Miller, M. P. Kent et al., 2016 The Atlantic salmon genome provides insights into rediploidization. Nature 533: 200–205.

Liu, L., K. P. Ang, J. A. K. Elliott, M. P. Kent, S. Lien et al., 2017 A genome scan for selection signatures comparing farmed Atlantic salmon with two wild populations: Testing colocalization among outlier markers, candidate genes, and quantitative trait loci for production traits. Evol. Appl. 10: 276–296.

Liu, J., H. Hu, S. Panserat, and L. Marandel, 2020 Evolutionary history of DNA methylation related genes in chordates: new insights from multiple whole genome duplications. Sci. Rep. 10: 1–14.

Liu, Y., K. D. Siegmund, P. W. Laird, and B. P. Berman, 2012 Bis-SNP: Combined DNA methylation and SNP calling for Bisulfite-seq data. Genome Biol. 13:.

Long, H. K., D. Sims, A. Heger, N. P. Blackledge, C. Kutter et al., 2013 Epigenetic conservation at gene regulatory elements revealed by non-methylated DNA profiling in seven vertebrates. Elife 2013: 1–19.

Le Luyer, J., M. Laporte, T. D. Beacham, K. H. Kaukinen, R. E. Withler et al., 2017 Parallel epigenetic modifications induced by hatchery rearing in a Pacific salmon. Proc. Natl. Acad. Sci. 114: 12964–12969.

McCormick, S. D., 2001 Endocrine control of osmoregulation in teleost fish. Am. Zool. 41: 781–794.

McNamara, J. M., S. R. X. Dall, P. Hammerstein, and O. Leimar, 2016 Detection vs. selection: integration of genetic, epigenetic and environmental cues in fluctuating environments (T. Coulson, Ed.). Ecol. Lett. 19: 1267–1276.

Metzger, D. C. H., and P. M. Schulte, 2017 Persistent and plastic effects of temperature on DNA methylation across the genome of threespine stickleback (*Gasterosteus aculeatus*). Proc. R. Soc. B Biol. Sci. 284: 20171667.

Monaghan, P., and N. B. Metcalfe, 2019 The deteriorating soma and the indispensable germline: gamete senescence and offspring fitness. Proc. R. Soc. B Biol. Sci. 286: 20192187.

Naish, K. A., J. E. Taylor, P. S. Levin, T. P. Quinn, J. R. Winton et al., 2007 An Evaluation of the Effects of Conservation and Fishery Enhancement Hatcheries on Wild Populations of Salmon, pp. 61–194 in Advances in Marine Biology,.

Nakamura, T., M. Komiya, K. Sone, E. Hirose, N. Gotoh et al., 2002 Grit, a GTPase-Activating Protein for the Rho Family, Regulates Neurite Extension through Association with the TrkA Receptor and N-Shc and CrkL/Crk Adapter Molecules. Mol. Cell. Biol. 22: 8721–8734.

O’Reilly, P., and R. Doyle, 2007 Live gene banking of endangered populations of Atlantic salmon, pp. 425–469 in The Atlantic Salmon: Genetics, Conservation and Management, edited by E. Verspoor, L. Stradmeyer, and J. Nielsen. Blackwell Publishing Ltd, Oxford, UK.

O’Rourke, T., and C. Boeckx, 2020 Glutamate receptors in domestication and modern human evolution. Neurosci. Biobehav. Rev. 108: 341–357.

O’Sullivan, R. J., T. Aykanat, S. E. Johnston, G. Rogan, R. Poole et al., 2020 Captive-bred Atlantic salmon released into the wild have fewer offspring than wild-bred fish and decrease population productivity. Proceedings. Biol. Sci. 287: 20201671.

Ohlsson, C., S. Mohan, K. Sjo□gren, A. Tivesten, J. Isgaard et al., 2009 The Role of Liver-Derived Insulin-Like Growth Factor-I. Endocr. Rev. 30: 494–535.

Oksanen, J., F. G. Blanchet, M. Friendly, R. Kindt, P. Legendre et al., 2019 vegan: Community Ecology Package. R Packag. version 2.5–6.

Ortega-Recalde, O., R. C. Day, N. J. Gemmell, and T. A. Hore, 2019 Zebrafish preserve global germline DNA methylation while sex-linked rDNA is amplified and demethylated during feminisation. Nat. Commun. 10: 1–10.

Paradis, E., 2010 pegas: an R package for population genetics with an integrated-modular approach. Bioinformatics 26: 419–420.

Pedersen, B. S., K. Eyring, S. De, I. V. Yang, and D. A. Schwartz, 2014 Fast and accurate alignment of long bisulfite-seq reads. arXiv.

Potok, M. E., D. A. Nix, T. J. Parnell, and B. R. Cairns, 2013 Reprogramming the Maternal Zebrafish Genome after Fertilization to Match the Paternal Methylation Pattern. Cell 153: 759–772.

Quinlan, A. R., and I. M. Hall, 2010 BEDTools: a flexible suite of utilities for comparing genomic features. Bioinformatics 26: 841–842.

Quinn, T. P., and N. P. Peterson, 1996 The influence of habitat comple×ity and fish size on overwinter survival and growth of individually marked juvenile coho salmon (*Oncorhynchus kisutch*) in Big Beef Creek, Washington. Can. J. Fish. Aquat. Sci. 53: 1555–1564.

R Core Team, 2019 A language and environment for statistical computing.

Richards, E. J., 2006 Inherited epigenetic variation — revisiting soft inheritance. Nat. Rev. Genet. 7: 395–401.

Rodriguez Barreto, D., C. Garcia de Leaniz, E. Verspoor, H. Sobolewska, M. Coulson et al., 2019 DNA Methylation Changes in the Sperm of Captive-Reared Fish: A Route to Epigenetic Introgression in Wild Populations. Mol. Biol. Evol. 36: 2205–2211.

Rougeux, C., P. Gagnaire, K. Praebel, O. Seehausen, and L. Bernatchez, 2019 Polygenic selection drives the evolution of convergent transcriptomic landscapes across continents within a Nearctic sister species complex. Mol. Ecol. 28: 4388–4403.

Ryan, D., 2019 MethylDackel. GitHub Repos.

Ryu, T., H. D. Veilleux, J. M. Donelson, P. L. Munday, and T. Ravasi, 2018 The epigenetic landscape of transgenerational acclimation to ocean warming. Nat. Clim. Chang. 8: 504–509.

Schaffer, A. E., M. W. Breuss, A. O. Caglayan, N. Al-Sanaa, H. Y. Al-Abdulwahed et al., 2018 Biallelic loss of human CTNNA2, encoding αN-catenin, leads to ARP2/3 complex overactivity and disordered cortical neuronal migration. Nat. Genet. 50: 1093–1101.

Sciamanna, I., A. Serafino, J. A. Shapiro, and C. Spadafora, 2019 The active role of spermatozoa in transgenerational inheritance. Proc. R. Soc. B Biol. Sci. 286: 20191263.

Shrimpton, J. M., N. J. Bernier, G. K. Iwama, and D. J. Randall, 1994 Differences in Measurements of Smolt Development Between Wild and Hatchery-Reared Juvenile Coho Salmon (*Oncorhynchus kisutch*) Before and After Saltwater Exposure. Can. J. Fish. Aquat. Sci. 51: 2170–2178.

Skaala, Ø., F. Besnier, R. Borgstrøm, B. T. Barlaup, A. G. Sørvik et al., 2019 An extensive common-garden study with domesticated and wild Atlantic salmon in the wild reveals impact on smolt production and shifts in fitness traits. Evol. Appl. 12: 1001–1016.

Skvortsova, K., N. Iovino, and O. Bogdanović, 2018 Functions and mechanisms of epigenetic inheritance in animals. Nat. Rev. Mol. Cell Biol. 19: 774–790.

Skvortsova, K., K. Tarbashevich, M. Stehling, R. Lister, M. Irimia et al., 2019 Retention of paternal DNA methylome in the developing zebrafish germline. Nat. Commun. 10: 1–13.

Stark, E. J., E. J. Atkinson, and C. C. Kozfkay, 2014 Captive rearing for Chinook salmon (*Oncorhynchus tshawytscha*) and Atlantic salmon (*Salmo salar):* the Idaho and Maine experiences. Rev. Fish Biol. Fish. 24: 849–880.

Sutherland, B. J. G., J. M. Prokkola, C. Audet, and L. Bernatchez, 2019 Sex-Specific Coexpression Networks and Sex-Biased Gene Expression in the Salmonid Brook Charr Salvelinus fontinalis. G3&#58; Genes|Genomes|Genetics 9: g3.200910.2018.

Thorstad, E. B., F. Whoriskey, A. H. Rikardsen, and K. Aarestrup, 2010 Aquatic Nomads: The Life and Migrations of the Atlantic Salmon, pp. 1–32 in Atlantic Salmon Ecology, edited by Ø. Aas, S. Einum, A. Klemetsen, and J. Skurdal. Wiley-Blackwell, Oxford, UK.

Trefts, E., M. Gannon, and D. H. Wasserman, 2017 The liver. Curr. Biol. 27: R1147–R1151.

Vandersteen, W. E., R. Leggatt, L. F. Sundström, and R. H. Devlin, 2019 Importance of Experimental Environmental Conditions in Estimating Risks and Associated Uncertainty of Transgenic Fish Prior to Entry into Nature. Sci. Rep. 9: 406.

Wang, X., and R. K. Bhandari, 2019 DNA methylation dynamics during epigenetic reprogramming of medaka embryo. Epigenetics 14: 611–622.

Wang, X., J. A. Weiner, S. Levi, A. M. Craig, A. Bradley et al., 2002 Gamma Protocadherins Are Required for Survival of Spinal Interneurons. Neuron 36: 843–854.

Whitlock, M. C., and K. E. Lotterhos, 2015 Reliable Detection of Loci Responsible for Local Adaptation: Inference of a Null Model through Trimming the Distribution of F ST. Am. Nat. 186: S24–S36.

